# 3D markerless tracking of speech movements with submillimeter accuracy

**DOI:** 10.1101/2025.02.13.638009

**Authors:** Austin Lovell, James Liu, Arielle Borovsky, Raymond A. Yeh, Kwang S. Kim

**Affiliations:** Department of Computer Science, Purdue University, West Lafayette, IN, United States; Purdue University Interdisciplinary Life Science Program, Purdue University, West Lafayette, IN, United States; Department of Speech, Language, and Hearing Sciences, Purdue University, West Lafayette, IN, United States

## Abstract

Speech movements are highly complex and require precise tuning of both spatial and timing of oral articulators to support intelligible communication. These properties also make measurement of speech movements challenging, often requiring extensive physical sensors placed around the mouth and face that are not easily tolerated by certain populations such as young children. Recent progress in machine learning-based markerless facial landmark tracking technology demonstrated its potential to provide lip tracking without the need for physical sensors, but whether such technology can provide submillimeter precision and accuracy in 3D remains unknown. Moreover, it is also unclear whether such technology can be applied to track speech movements in young children. Here, we developed a novel approach that integrates Shape Preserving Facial Landmarks with Graph Attention Networks (SPIGA), a facial landmark detector, and CoTracker, a transformer-based neural network model that jointly tracks dense points across a video sequence. We further examined and validated this novel approach by assessing its tracking precision and accuracy. The findings revealed that our approach that integrates SPIGA and CoTracker was more precise (≈ 0.15 mm in standard deviation) than SPIGA alone (≈ 0.35 mm). In addition, its 3D tracking performance was comparable to electromagnetic articulography (≈ 0.29 mm RMSE against simultaneously recorded articulograph data). Importantly, the approach performed similarly well across adults and young children (*i*.*e*., 3- and 4-year-olds). Because our framework is built upon open-source pretrained models that are fully trained, it promotes accessibility and open science while saving computing resources. Furthermore, given that this framework combines a landmark detection model (SPIGA) with a tracker model (CoTracker) to improve precision/accuracy, our novel approach serves as a proof-of-concept for enhancing the performance of a wide variety of commonly used markerless tracking applications in biology and neuroscience.

**Author summary:** In this work, we examined whether machine learning based markerless tracking is feasible for tracking 3D lip movements in adults and young children. We developed a novel approach that integrates a landmark detection model (SPIGA) with a tracker model (CoTracker). Our combined CoTracker-based approach demonstrated submillimeter precision and accuracy desired for speech kinematic recording. In addition, our approach does not involve training and validation for each population (*e*.*g*., young children vs. adults), saving time and computing resources. We foresee that the proposed general framework of fusing a landmark detection model with a tracker model can be generalized for a wide variety of tracking applications in biology and neuroscience that require high precision and accuracy, including studying cell behaviors, animal motions, and other types of human movements.

## Introduction

Recording speech movements is critical for investigating speech motor control in individuals without any speech disorders (*e*.*g*., adults, [1, 2], children, [3, 4]) as well as those with speech and language difficulties (*e*.*g*., developmental language disorder [5], stuttering [6–8], Parkinson’s disease [9], and amyotrophic lateral sclerosis [10]). To date, multiple methods have been applied to track speech movements (see Earnest & Max [11] for a detailed review). For example, a strain gauge system that makes use of cantilever beam transduction (*i*.*e*., metal strips bending due to speech movements) has been used for multiple decades, *e*.*g*., Barlow & colleagues [12, 13]. Other well-known methods include X-ray microbeam (*e*.*g*., Westbury [14]) and ultrasound (*e*.*g*., Stone & Lundberg [15] Whalen et al. [16]) imaging methods. In addition, electromagnetic articulography, which captures articulatory movements by generating an electromagnetic field and tracking locations of sensor coils, has also been widely used (*e*.*g*., Yunusova et al. [17]). Researchers have also used optical motion capture systems that track infrared light-emitting markers (*e*.*g*., Vuolo & Goffman [18]), and custom video-based systems that track reflective markers [19].

However, available methods are not ideal for young children (*i*.*e*., age 4 or below) who typically do not tolerate physical sensors and markers. We seek to overcome this challenge by building on recent advances in the ability of machine learning models to track movements in humans [20, 21] and animals [22] from video recordings—without any physical sensors or markers. For example, machine learning models have been able to perform comparably to human labeling in tracking lip movements (and tongue movements based on ultrasound images) through software such as DeepLabCut, a well-established pose estimation method based on deep neural networks [23].

Markerless tracking is promising because it not only eliminates the need for sensors and calibration/preparation steps required for the sensors but is also more affordable than commercially available speech tracking systems [23]. In addition, it is an accessible method that promotes open science given that many models and software are usually open-source. Furthermore, the approach also allows for ecologically-valid measures of natural movement “outside the lab” given that video recording equipment is highly portable.

Although the facial landmark detection abilities of currently available machine learning models are considered to be sufficient in the field of computer vision [24–27], their precision and accuracy for speech movement tracking applications remain largely unknown. Because speech articulator movements can be as small as a few millimeters, it is important to achieve submillimeter accuracy and precision [19]. Furthermore, it also remains unknown whether these types of technology can be applied to track speech movements in young children. Currently, publicly available datasets that are used to train facial landmark detection tend to under-represent children, focusing primarily on adults [26, 28, 29]. Because children’s faces differ from those of adults (*e*.*g*., the spatial proportions of facial features are very different), a neural network model trained solely on adults’ facial landmarks may not reliably track speech movements in young children.

Overcoming these challenges is not trivial, given that training machine learning models that can generalize robustly across individuals and populations is time and resource consuming.

Here, we developed a novel approach by leveraging a pretrained model to detect facial landmarks (including the lips) in the initial frames of the video clips and applying another model to track multiple landmarks of interest across the frames in videos. Specifically, Shape Preserving Facial Landmarks with Graph Attention Networks (SPIGA [27]), a markerless face alignment and head pose estimator, performed initial facial feature detection, and CoTracker [30], a transformer-based neural network model, tracked the points of interest jointly across a video sequence based on the coordinates provided by SPIGA in the initial frames. As an initial step towards validating this combined approach for lip tracking, we assessed whether it provides precise and accurate lip aperture (LA) measurements in 3D (*i*.*e*., the Euclidean distance between the upper lip and lower lip). First, we examined whether CoTracker’s lip aperture measures of static lips remained constant despite head movements. Second, tracking accuracy was quantified by comparing CoTracker’s LA measures against measurements obtained by an electromagnetic articulograph (AG501, Carstens), one of the most reliable and accurate electromagnetic systems [31]. Importantly, lip tracking was performed in both adults and young children to test the feasibility of this combined approach for a diverse population.

## Approach Overview

### Preprocessing

To obtain 3D kinematic data, two GoPro cameras were placed approximately 80 cm in front of participants, spaced about 60 cm from each other. Using the cameras, we obtained video recordings of participants making bilabial utterances such as “Buy Bobby a puppy” (adults) or “puppy” (children). These videos were time-synchronized using a clapboard sound (see “Time Sync Stereo Videos” in Fig 1) and calibrated using OpenCV [32] (see Materials and Methods for more details). Video recordings (5.3K resolution, 60 Hz, see Materials and Methods for more details) were then trimmed into shorter clips for each trial or utterance (*i*.*e*., 0.5 s to 3 s in this study) using FFmpeg [33].

**Fig 1.**
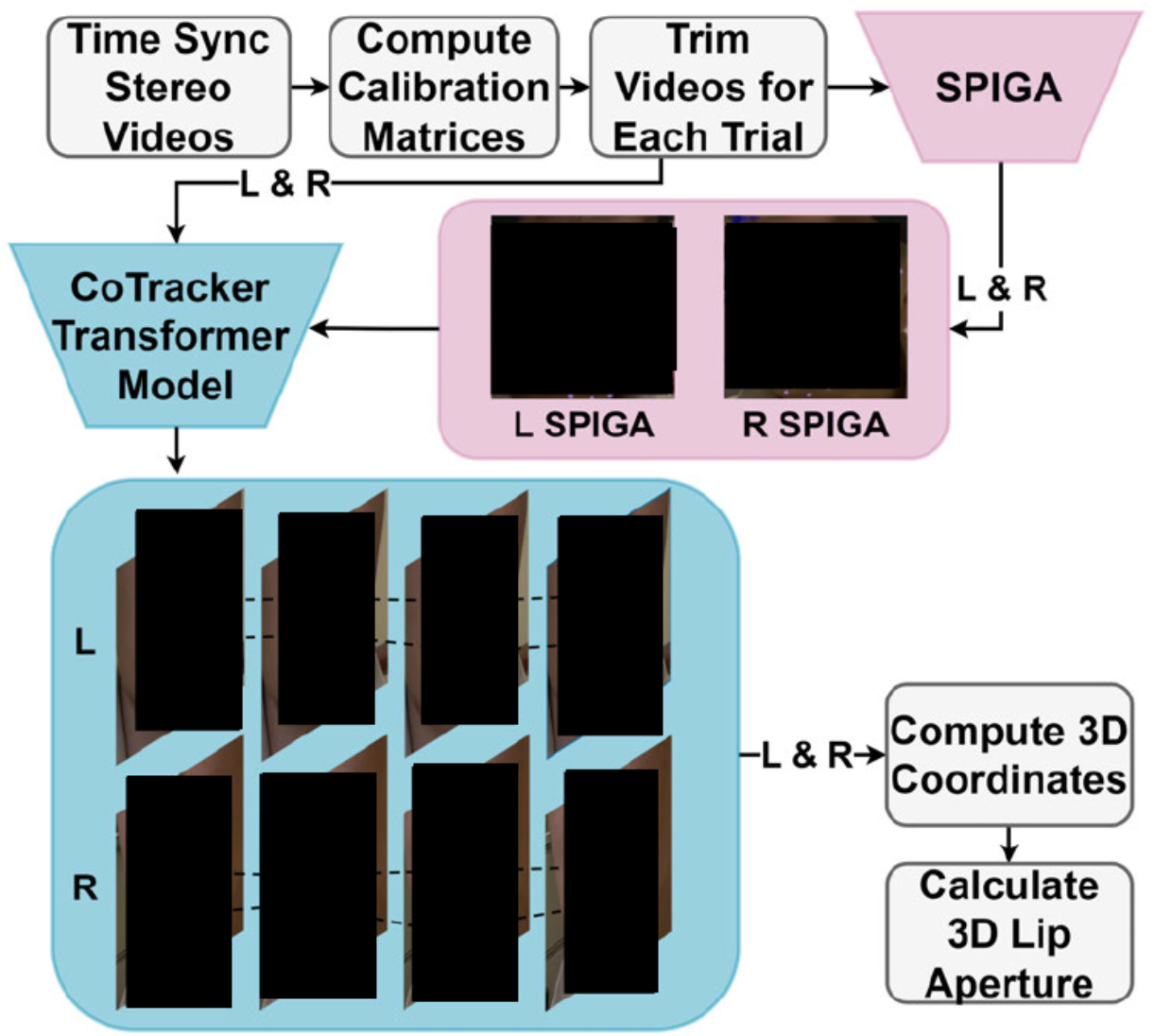
CoTracker Pipeline. After data collection with two GoPro cameras, the videos (i.e., “L” for the left video and “R” for the right video in the perspective of the experimenters facing the participant) were time-synchronized using a clapboard sound (“Time Sync Stereo Videos”). Ea.ch video included 5 to 10 seconds of calibration checkerboard motions, which were used for calibrating using the OpenCV library [32] (“Compute Calibration Matrices”). After the videos were trimmed for each trial using FF mpeg [33], SPIGA placed points on facial landmarks on the initial frames for each video (see pink color points). Using the initial lip coordinates, CoTracker tracked the lip points of interest (see light blue points). The pixel coordinates from each video were used to compute 3D coordinates in millimeters (“Compute 3D Coordinates”). The 3D lip aperture data was then calculated.

### SPIGA

Shape Preserving Facial Landmarks with Graph Attention Networks (SPIGA, see Fig 1) is a neural network model that estimates facial landmarks in images or videos [27]. Specifically, SPIGA utilizes traditional convolutional neural networks (CNN) in combination with a cascade of Graph Attention Network (GAT) regressors. This algorithm provides robust performance in different contexts including occlusion, heavy makeup, blur, and extreme illumination. For our pipeline, SPIGA was utilized to estimate the initial starting points of the desired lip locations as well as other facial landmarks to serve as “supporting” points for CoTracker (see CoTracker for more details). The ten initial landmark points on the lip were determined by taking the average of SPIGA’s coordinates for these points across the first five frames of a video clip.

### CoTracker

CoTracker [30] is a transformer-based model designed to track points (or coordinates) through a video sequence. It is capable of tracking dense points, allowing users to create grids of points across an entire video to track as compared to other models that can only process a few individual points. Importantly, the CoTracker model uses joint tracking, which means that, for a specific point, it correlates information between other tracked points in a sequence to improve prediction efficacy. This design is drastically different from alternative models that were constrained to independent tracking where each point is tracked individually [34, 35].

In our pipeline, CoTracker used initial coordinate information provided by SPIGA on the first five frames of the videos. CoTracker tracked the specified points across the frames for both the left and right videos.

### Postprocessing

After the points of interest were determined in terms of pixel coordinates by CoTracker, OpenCV library methods [32] were used to convert 2D left and right video pixel coordinates to 3D coordinates for each point in a real-world unit (*i*.*e*. millimeter). We then calculated lip aperture, the Euclidean distance between the mid-sagittal upper and lower vermilion border in the 3D space (*e*.*g*., Benham et al. [5] or Walsh et al. [7], and also see Methods for more details).

## Results

### Lip aperture precision during head movements

As the first validation test, we examined whether 3D lip aperture (LA) measurements of static lips (*i*.*e*., not moving lips) remained constant during head movements. We analyzed tracking precision by calculating the standard deviation of lip aperture across frames. Specifically, we compared CoTracker’s performance with that of SPIGA, because SPIGA, unlike CoTracker, does not correlate information across frames (*i*.*e*., SPIGA identifies landmarks on each frame independently). Thus, this comparison serves as an ablation analysis that determines whether CoTracker is a critical component in our pipeline for consistent and precise tracking. It should be noted that in this section we refer to our CoTracker pipeline in which SPIGA provides facial landmarks for the initial frames but CoTracker does the actual tracking as “CoTracker,” in order to distinguish it from the SPIGA-only pipeline (“SPIGA”).

We obtained video recordings from three children (age M = 37.0, SD = 0.5 months) whose data was recorded as part of another study. During the sessions, we identified brief periods during which participants did not move their lips (*i*.*e*., static lip aperture), but moved their heads to a small extent. During these moments, head movements resulted from participants slightly adjusting their posture or changing their head orientation as they were listening to the experimenters or waiting for certain cues in between speech tasks (“minimal head movement” see Fig 2 top row). These moments were trimmed into short, one-second clips, but some moments that lasted shorter than 1 second were trimmed into 0.5 second long clips instead (22 out of 123 clips).

**Fig 2.**
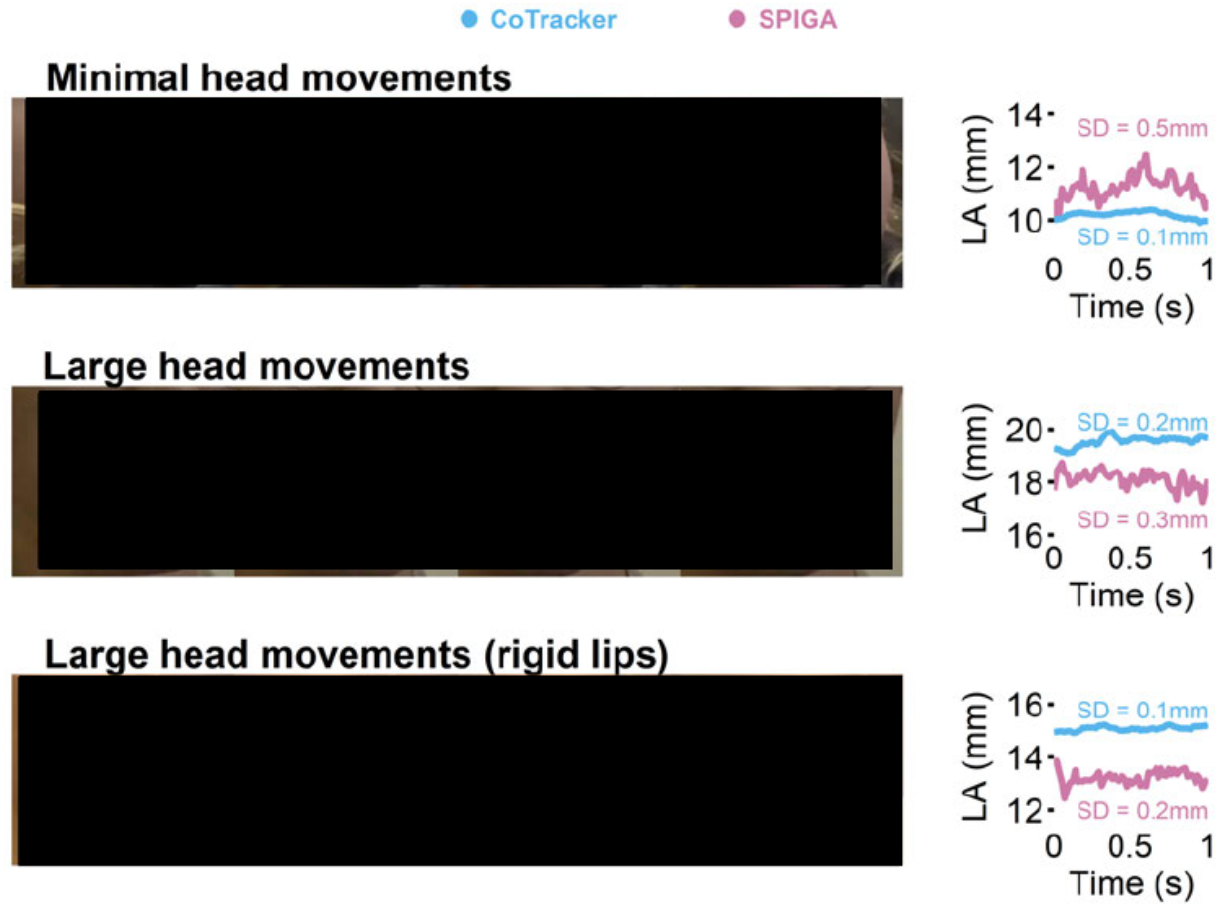
Static lip aperture tracking by CoTracker and SPIGA. We obtained videos of participants slightly moving their heads while their lips were relatively still (top left). Some participants were asked to make large head movements while not moving their lips (middle left). In a.ddition, a rigid mannequin head was placed on a rotating display (bottom left). Points placed by CoTracker (light blue) and SPIGA (pink) are shown on these panels. The right panels show the 3D lip aperture **(LA)** computed from the points and the standard deviation of **LA**. The ideal result is a low standard deviation (SD), with a flat, horizontal **LA** across time, which would suggest a high degree of precision. It should be noted that in some cases CoTracker’s starting LA amplitudes were slightly different from SPIGA’s, because an edge detection-based adjustment was performed to place the initial landmarks more closely to vermilion borders of the lips for CoTracker (see Materials and Methods for more details).

To examine lip aperture precision in a more challenging context, we asked 5 adult participants (age M = 28.0, SD = 6.6 years) and 2 child participants (age M = 43.0, SD = 1.4 months) to make larger head movements by alternating their head orientation (*i*.*e*. looking at each camera one after the other, see “large head movement” in Fig 2 middle row). For some adult participants, we also asked them to move their heads up and down during these movements. We also considered the possibility that participants could have been making extremely small lip movements in these videos. Therefore, to further test the stability and robustness of our lip tracking technology against head movements, we also sought to test a rigid artificial head model, where no lip movement could be assured. Thus, we placed a rigid mannequin head on a rotating display stand that rotates 18 degrees per second in the transverse plane (see Fig 2 bottom row) to further assess whether and to what degree the model inaccurately registered lip movement in the presence of head movements.

We analyzed these stationary lip videos in the presence of head movement in a total of 132 video clips from 10 participants (5 children, 5 adults) and 1 rigid head model. LA was extracted from 1 second-long videos (*i*.*e*., 60 frames) in most cases, but we also extracted LA from shorter moments (500 ms) in some trials due to participants moving their lips or other tracking issues (*e*.*g*., the camera view is occluded by a hand). The standard deviation of the LA measures was calculated to assess the tracking precision of CoTracker and SPIGA (see the right panels in Fig 2).

Our measures indicated that both models exhibited sub-millimeter precision in LA artifacts in the presence of head movement. The overall average standard deviations of LA for CoTracker and SPIGA were 0.104 mm and 0.379 mm respectively (see Fig 3). A linear mixed effects model revealed that LA standard deviation was significantly lower in CoTracker compared to SPIGA during minimal head movements, F(1, 62.244) =262.782, p < 0.001. The fixed effect of the length of the video clips (i.e. 0.5 s vs. 1 s) was not significant, F(l, 64.439) = 0.392, p = 0.533. There was a significant interaction between the video clip length and tracking method (i.e. CoTracker vs. SPIGA), F(l, 62.244) = 4.029, p = 0.049. A post hoc test of pairwise comparison revealed that the only significantly different comparisons were found in the pairs with the different tracking methods.

**Fig 3.**
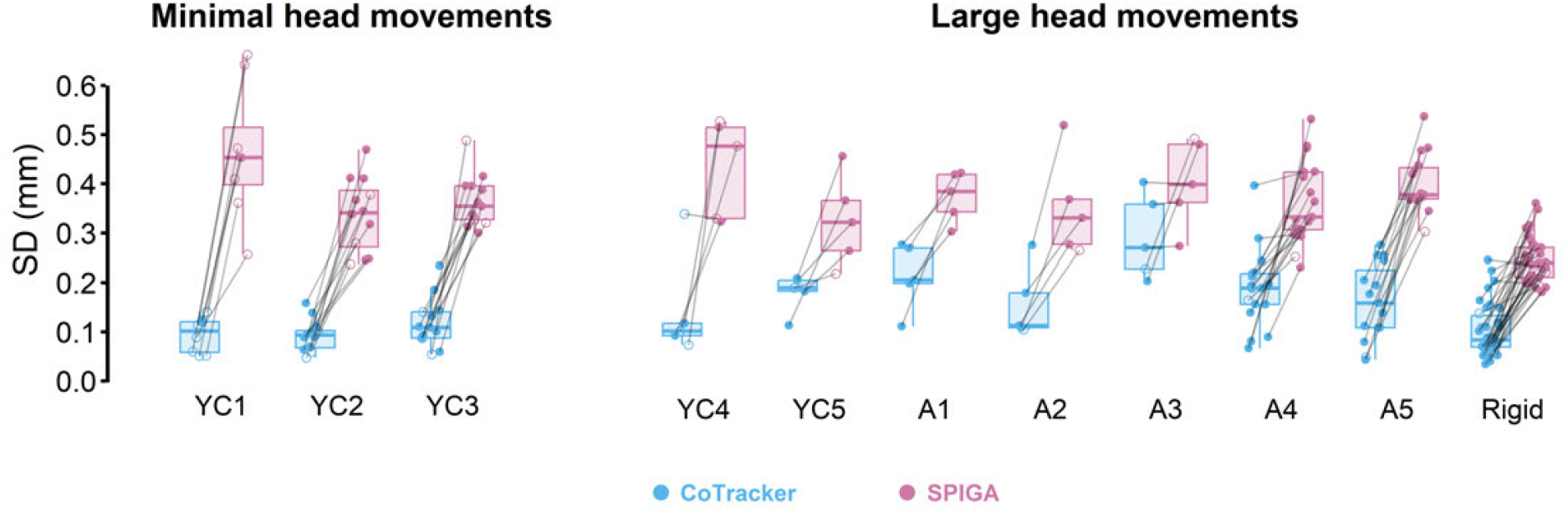
Standard deviation of the lip aperture measures in each participant. Each point represents standard deviation in a given video clip. The points are connected with gray lines to denote that tracking yielded from the same video clips. Overall, CoTracker (light blue) provided more consistent tracking (*i*.*e*. lower SD) than the SPIGA-only pipeline (pink) across all participants including young children (labeled as “YC”) and adults (labeled as “A”). We also found the same pattern from mannequin lips (labeled as “Rigid”). We analyzed a total of 132 video clips (110 videos that were 1 second-long shown as solid circles and 22 clips that were 0.5 second-long shown as transparent circles.

The standard deviations during large head movements, 0.160 mm (CoTracker) and 0.334 mm (SPIGA), were comparable to the overall average, suggesting that the precision was not affected by larger movements. A separate linear mixed effects model for larger head movements data demonstrated that, again, CoTracker provided more consistent tracking (*i*.*e*., significantly lower standard deviation) than SPIGA did, F(1, 174.83) = 274.23, p < 0.001. On the other hand, the fixed effect of the video length was not significant, F(1, 181.39) = 3.292, p = 0.071. The interaction between the tracking method and the video length was also not significant, F(1, 174.69) = 0.073, p = 0.787.

In sum, across both minimal and large head movements, CoTracker performed with a higher precision (*i*.*e*., average 0.145 mm SD across all trials) than SPIGA alone (*i*.*e*. 0.346 mm). Our findings add support to our framework in which SPIGA provides initial landmark and CoTracker (as opposed to SPIGA) tracks them throughout the video clips. Notably, the standard deviation remained low for young children as well (see “YC” in Fig 3). The standard deviation for CoTracker was larger than 0.3 mm only in 4 trials (of the 132 trials). Together, these findings clearly suggest that CoTracker offers submillimeter precision, suitable for tracking speech kinematic data which may be as small as a few millimeters [19].

### Lip aperture accuracy during speech production

As another validation test, we compared CoTracker with an electromagnentic articulograph (AG501, Carstens). In this test, LA measured by AG501 was used as the “ground truth,” given that it is arguably the most accurate speech kinematic recording system, offering mean error less than 0.2 mm [36]. Prior to recording, we attached two articulograph sensors on the participant’s upper and lower lips, at the mid-sagittal section of the vermilion borders (see Fig 4 for example), which is standard practice for measurement of LA (*e*.*g*., Vuolo & Goffman [18]). Participants were asked to sit in the articulograph that is placed in front of the cameras. For this test, we recruited 7 adults (age M = 24.9, SD = 6.1 years) and 3 children (age M = 45.3, SD = 4.2 months). We asked adult participants to repeat the phrase “Buy Bobby a puppy” 10 to 15 times.

**Fig 4.**
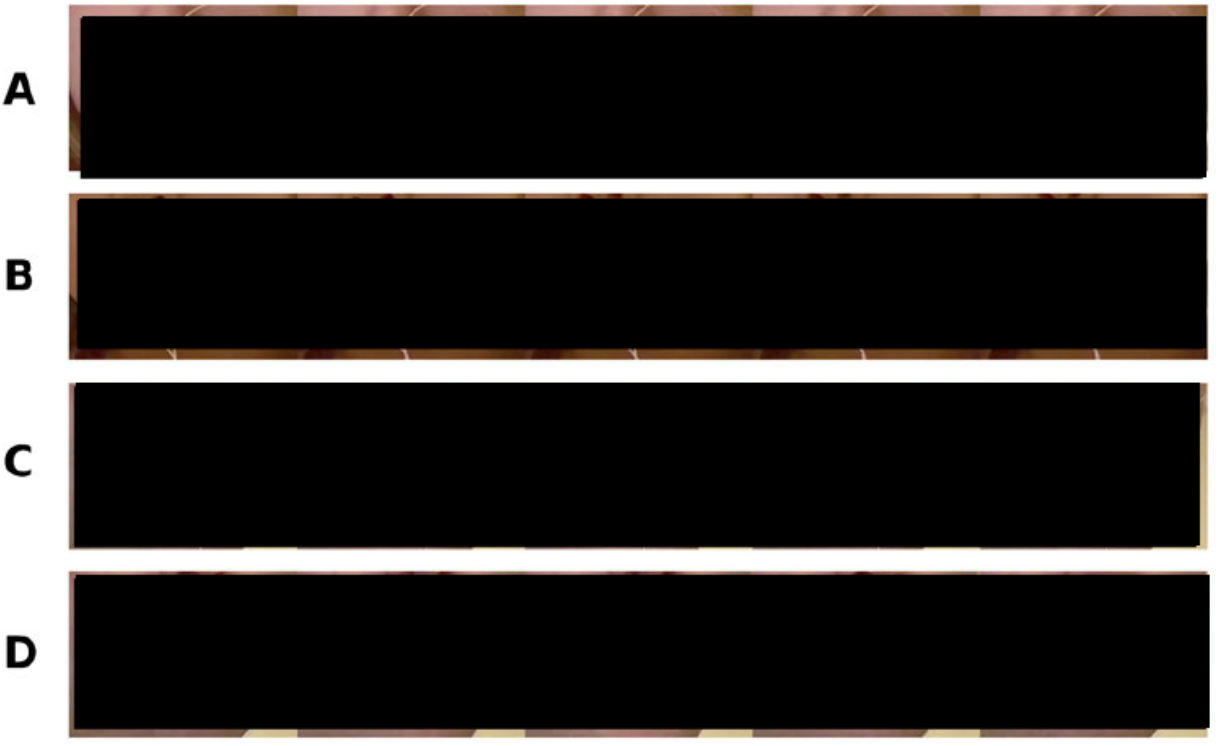
Simultaneous re cording of AG501 and video recordings. Participants were seated in the articulograph (AG501) with two sensors (black sensors with white wires) attached to their lips (one on the upper lip and the other on the lower lip). Two GoPro cameras were placed in front of the articulograph to record videos, which were processed with our CoTracker pipeline. In the top two rows (A, B), videos of adult and child participants are shown respectively. For a few participants, we were also able to remove AG501 sensors from the original videos for further validation tests of CoTracker (see the original video in C and the inpainted video in D). CoTracker’s points are shown in a light blue color in unaltered videos (A, B, and C) and green in the inpainted video (D). The brightness of the images was slightly enhanced to improve visualization in the figure.

This is a commonly used phrase in speech motor control research as it includes many bilabial phonemes that elicit many opportunities for changes in lip aperture in a short period of time (*e*.*g*., Smith & Goffman [37]). For the child participants, we asked them to say “puppy” following prior practice in the field, as it is possible for typically-developing young children of this age to remember and say, while also ensuring many opportunities to record bilabial movement *(e*.*g*., Vuolo & Goffman [181).

To compare lip aperture data between AG501 and CoTracker, we first upsampled CoTracker’s data to match AG501 sampling rate (*i*.*e*. 250 Hz). The upsampled CoTracker data were then time-synchronized with AG501 data by selecting a temporal alignment that would provide the lowest root mean square error (RMSE, see Materials and methods) against the AG501 data (see left panels in Fig 5). After this temporal alignment step, we selected the lip aperture during the utterances (see the shaded area in yellow in the right panels of Fig 5) and normalized the CoTracker’s data to have the same average as AG501 LA for each utterance. This amplitude-normalization step was necessary because the locations of articulograph sensors are not identical to the locations of the points placed by SPIGA (as part of the CoTracker pipeline). After normalizing the amplitudes of LA, RMSE was computed during those utterances (see Methods for more details).

**Fig 5.**
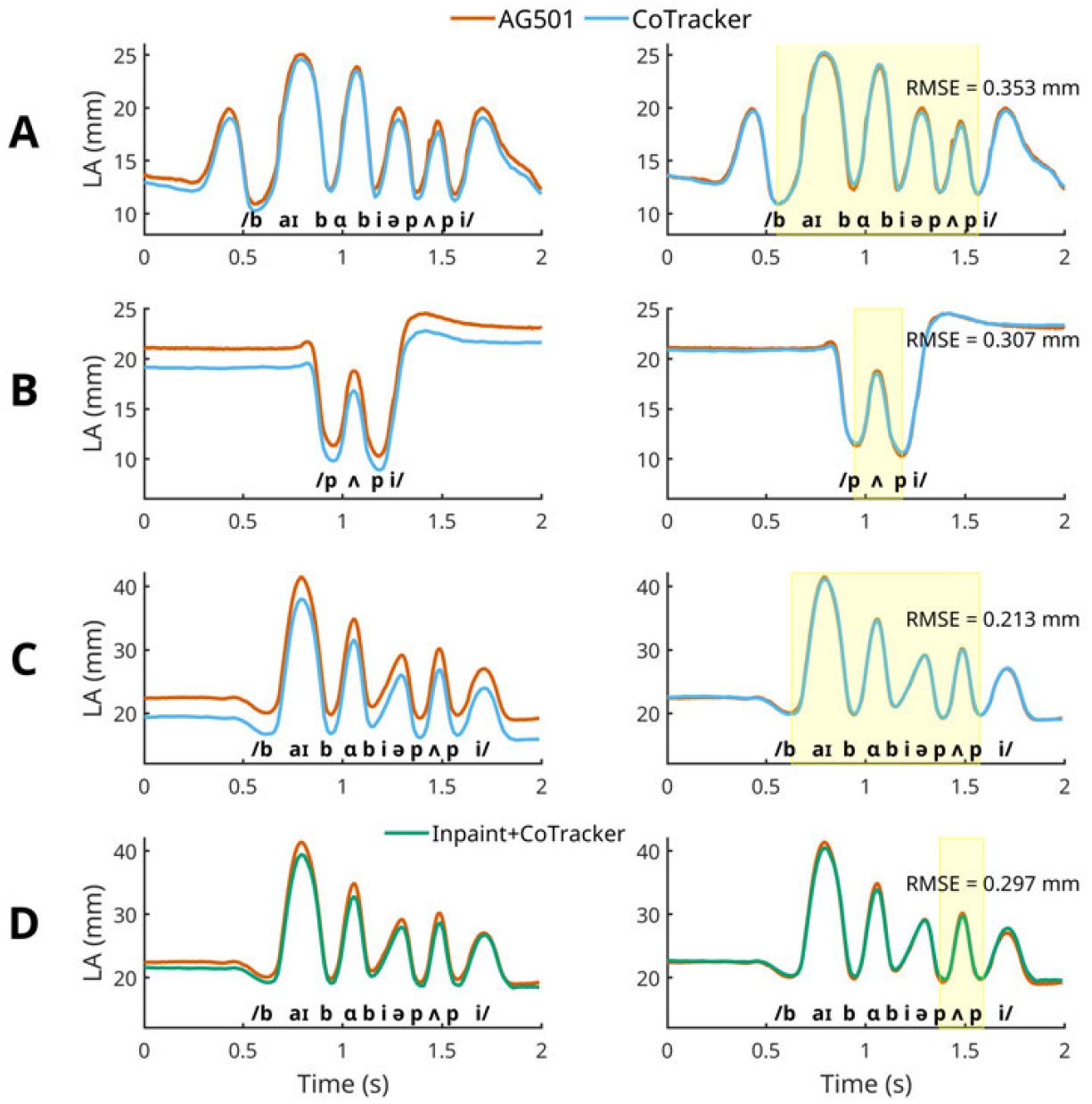
Lip Aperture data from AG501 vs. CoTracker. Data shown in each row of panels (A, B, C, and D) were generated from the corresponding videos (A, B, C, and D respectively) in Fig 4. International Phonetics Association transcription for the corresponding speech sounds is shown at the bottom of each panel. The left panels show time-synced but not amplitude-normalized CoTracker data. The right panels show time-synced and amplitude-normalized CoTracker data. The yellow shaded area denotes the frames during which utterances occurred. The amplitude-normalization was accomplished using the average values of each method (CoTracker vs. AG501) in these frames.

A total of 103 utterances from 10 different participants were analyzed (Fig 6). The results indicated that in both adult and child participants, the RMSE values were mostly lower than 0.5 mm (Fig 6). On average, RMSE was 0.290 mm. The average RMSE value is slightly larger than the average standard deviation (approximately 0.15 mm) found by the static lip tracking validation test, suggesting that tracking accuracy is slightly lower during speech production. Nonetheless, the median RMSE for all 10 participants was below 0.4 mm, demonstrating that CoTracker is capable of providing a submillimeter accuracy suitable for speech movement tracking [19].

**Fig 6.**
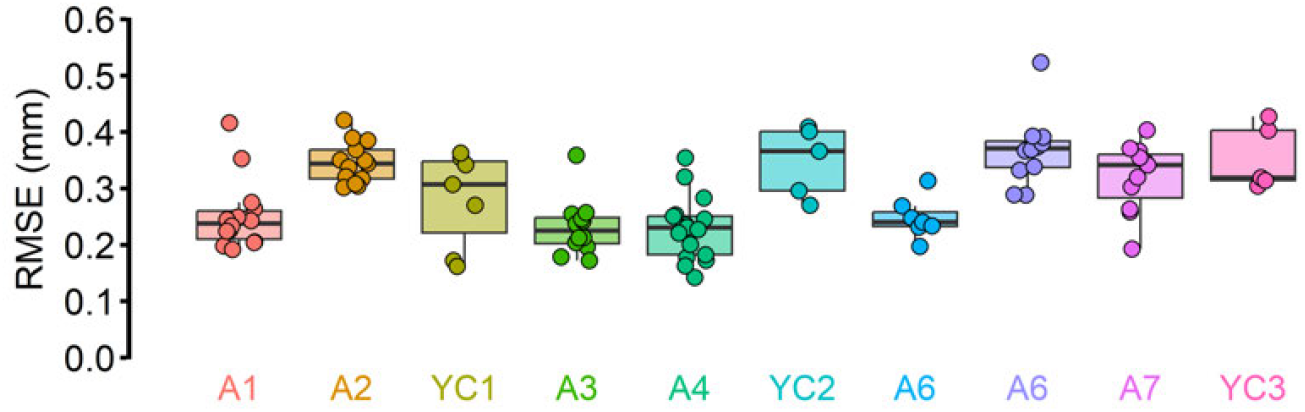
RMSE against AG501 across 10 participants. In both young children (“YC”) and adults (“A”), CoTracker’s tracking demonstrated a submillimeter accuracy when considering AG501 data as a “ground truth.” Although there was some degree of inter-individual variability, most trials (102 out of 103) had less than 0.5 mm RMSE. The median RMSE for all 10 participants was less than 0.4 mm, and the average RMSE across all participants was 0.290 mm.

### CoTracker in inpainted videos

As the articulograph sensors moved along with the lips, the presence of the sensors may have served as a “clue” for CoTracker to track lip landmarks. In other words, CoTracker’s tracking performance could have been enhanced by having the articulograph sensors attached. Hence, we conducted an additional test to rule out the possibility that CoTracker’s performance benefited from the presence of the sensors. A state-of-the-art machine learning inpainting algorithm was used to remove the articulograph sensors from each participant’s recording (Fig 4D).

First, a segmentation mask covering the articulograph sensors and their wires was created by Meta’s open-source Segment Anything 2 (SAM2 [38]) model for each frame of the video. The binary mask created by SAM2 (*i*.*e*., value 1 on pixels that cover the sensors) was then given as an input to an inpainting pipeline built on a pretrained open-source Hugging Face stable diffusion inpainting model (stable-diffusion-v1-5 [39]). The stable diffusion inpainting model was implemented to fill in the masked area with realistic, synthetic inpainted pixels. The 512 *×* 512 videos containing the covered synthetic data were then stitched back onto the original video for CoTracker. All inpainted videos followed the exact same time-sync and amplitude-normalization steps that were done for the other non-inpainted videos. It should be noted that because the inpainting quality was noticeably poor during larger movements such as “buy”, we decided to use a portion of the data (“pup”) from the inpainted videos. This is not surprising given that changes in the lips’ shape during large movements are likely to make the inpainting process more challenging.

The RMSEs for inpainted videos were mostly between 0.2 and 0.4 mm, similar to CoTracker’s performance on videos with the AG501 sensors (Fig 7). Although this analysis was performed with a small sample size (n = 3), this finding is consistent with our previous test in which CoTracker showed strong performance even when there were no physical sensors. Together, our results suggest that CoTracker’s performance was not enhanced by the presence of AG501 wires and that it can perform with a submillimeter accuracy even without physical sensors or visual markers.

**Fig 7.**
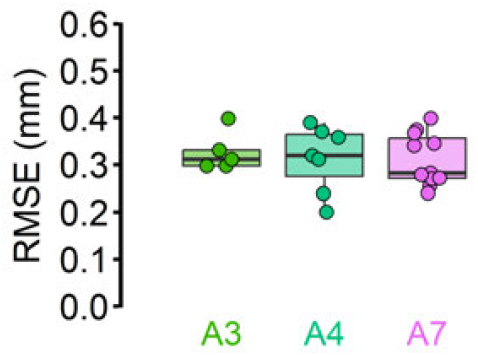
CoTracker on inpainted videos. Inpainted videos showed similar RMSE performance against AG501

## Discussion

We developed a novel markerless lip tracking approach combining SPIGA, a neural network-based facial landmark estimation model, and CoTracker, a transformer-based algorithm that tracks multiple points of interest by integrating information in a sequence of frames. Because this technology has a high potential for application to populations that would normally find traditional facial-marker based solutions challenging, such as young children, we sought to validate this approach in both young children and adults.

Our primary goal was to ensure that precise recording of lip movements could be accomplished with as few physical constraints as possible. Therefore, our first validation test sought to determine whether our solution had low lip movement artifact under conditions of large and small head movements. Here, we measured whether CoTracker’s lip aperture measures remain constant for static lips during head movements. This test demonstrated that the lip aperture measure was highly consistent in the presence of head movement (average standard deviation of 0.15 mm), suggesting that this approach has the potential to be used under conditions where participants’ natural movements are not restricted - an important advantage for developmental populations and those with communication challenges compared to current marker-based approaches.

Our second goal was to determine whether this approach can accurately measure lip movements at levels on par with current state-of-the-art marker-based technologies. To achieve this goal, we compared CoTracker’s lip aperture measures during speech production with those of an articulograph, and we found that the root mean square error (RMSE) against AG501 was about 0.290 mm on average. In both tests, CoTracker’s performance was comparable between young children and adults.

Given that both the static lip standard deviation and the RMSE against AG501 fall below 0.5 mm, it is clear that our markerless CoTracker system provides a high degree of accuracy in the measurement of lip movements, with measures that are very close to that of high-end marker-based systems. Importantly, given that lip aperture computation requires two coordinates, it is likely that each point’s precision/accuracy is smaller than those numbers reported in this investigation. These findings are somewhat surprising given that we used videos recorded by consumer-grade video cameras that had relatively low levels of spatial and temporal video resolution. It should also be noted that the degree of accuracy during movements in our markerless system is similar to other methods, *e*.*g*., RMSE around 0.26 mm in a custom camera-based system with reflective markers [19] or average Bland-Altman [40] error of 0.2 mm in AG501 [36].

Moreover, our approach is highly accessible for users who do not have access to high-performance computing resources. Even though our approach utilizes machine learning algorithms, this approach does not require end-user training or model tuning because it leverages well-established, pretrained models. For this reason, our combined approach does not require excessive graphic memory or high-end computing resources that are expensive and not widely available in consumer-grade workstations. In our pilot testing of video clips of different lengths, even 10 second long videos only required approximately 8.9 GB of GPU memory (including the baseline load required for the computer, see supplemental information I), which is well within the range of common consumer-grade graphic cards. Thus, consumer-grade computing equipment would be sufficient for longer utterances such as a sentence that lasts 10 seconds (see *e*.*g*., Darling-White & Banks [41]). In sum, our combined tracking approach is an accessible and affordable option for recording lip kinematics.

Given that our combined approach is novel and serves as an initial validation test, more validation tests are warranted. One limitation of the current study is that the dataset included only one participant with communication difficulties (*i*.*e*. language delay, see Materials and Methods). To increase our approach’s robustness, future work should examine tracking performance in larger sample sizes as well as a wide variety of clinical populations such as children who stutter.

In addition to increasing human participant datasets, future work should also incorporate other methods to serve as “ground truth” benchmarks. In the current study, AG501 was the only benchmark. Even though AG501 is thought to be superior compared to other electromagnetic systems [31, 42], it does not always provide accurate measures [36]. One alternative benchmark may be provided by hand labeling as demonstrated by Wrench & Balch-Tomes [23]. Another potential reference may be generated by synthetic images or videos created by machine learning techniques.

Future studies should also investigate the effect of lighting on tracking performance. It is intuitive that computer vision tasks, such as markerless tracking, require an adequate amount of lighting. However, we also observed that too much lighting created reflections that make markerless tracking challenging, likely because the reflections of the light source on the lips do not move along with the lips. Hence, it is important to determine the optimal intensity and directionality of lighting for lip tracking.

One limitation in our approach is that SPIGA, which provided the initial facial landmarks for CoTracker, is not precise (see Fig 3). These imprecise marker coordinates may lead to inconsistency in lip aperture measures across different trials (*i*.*e*. across different video clips). The main cause of this limitation is that SPIGA was trained to detect facial features on 256 *×* 256 pixel size images that contain faces. Thus, even high-resolution images are first downsampled to 256 *×* 256 pixel size images when facial landmarks are identified, which leads to the facial landmarks’ poorer resolution.

In the current study, SPIGA’s lack of accuracy was not a major issue given that each trial’s lip aperture was normalized in the accuracy test (see Fig 5). Additionally, it should be noted that for some analyses of lip aperture variability such as spatiotemporal index, SPIGA’s inter-trial variability does not pose a serious problem because the analysis is based on normalized LA amplitude across multiple productions (*e*.*g*., Vuolo & Goffman [18]). Nonetheless, if the use of the non-normalized data is necessary, SPIGA’s performance may be a bottleneck for tracking accuracy and will increase inter-trial variability. One potential solution that future studies should explore is whether it is possible to improve SPIGA’s accuracy by further training SPIGA with new datasets of high-resolution images with more accurate annotations of the vermilion borders of the lips.

## Conclusion & future directions

Our novel approach is capable of providing submillimeter precision and accuracy for 3D lip movement tracking in both adults and young children. Given the continual and rapid progress in artificial intelligence, our approach has a high potential for further improvement in near future. In addition, our approach may be translated into future clinical applications, such as providing a more accessible assessment of speech motor disorders. Furthermore, this proposed general framework of fusing a landmark detection model with a tracker model may be generalized for a wide variety of tracking applications in biology and neuroscience that require high precision and accuracy, including studying cell behaviors, animal motions, and other types of human movements.

## Materials and methods

### Participants

Eleven adults (age M = 26.2, SD = 6.55 years) and seven children (age M = 41.3, SD = 5.1 months) participated in at least one task (*i*.*e*. precision or accuracy tests) in the study. All adult participants had no known speech, language, or hearing disorders. One child had a language delay (*<*87 standard score on the SPELT [43]). Two of the children participated in both tests. All participants provided written informed consent (adults, and guardians or parents of the child participants) or verbal assent (children).

### Data collection

We placed two GoPro cameras (Hero 11 Black, GoPro) about 80 cm apart from the participant’s chair. The two cameras were also placed about 60 cm from each other to generate 3D tracking via triangulation. The videos were recorded with 5.3K resolution (5312 × 2988) at 60 Hz, using the Linear lens setting. To synchronize the timing of the videos, we generated a clap sound via a clapboard after the recording started. We also waved a commercially available checkerboard with 18 rows by 25 columns of squares with 1.5 cm sides (calib.io, Svendborg, Denmark) for about 5-10 seconds to calibrate cameras for 3D estimation. After these two steps, participants were seated in front of the cameras.

It should be noted that SPIGA is designed to label all faces it can detect in a given video. Thus, on some occasions, when faces other than the subjects’ were present in the video, SPIGA would track those other faces. To prevent this confusion, a Python script was used to “censor” unwanted faces with a black rectangle throughout the video. In addition, we excluded some trials due to occlusion (*i*.*e*., AG501 sensor wires occluding points of interest or child participants turning their heads to a degree in which the mid-sagittal portions of the lips are not visible).

### Software design

All codes used in our pipeline were written in Python (version 3.10.12, [44]) and are available on a GitHub repository (https://github.com/austinlovell25/cotracker).

#### Preprocessing

We first determined the timing of the sound of the clapboard by searching for the maximum peaks in the audio track of each video based on the user-defined threshold. Using these peaks (*i*.*e*., clap sound), we synchronized the times of the two videos (see Fig 1). In addition, from each video, we extracted frames that contain the checkerboard to calibrate each camera and randomly chose 50 frames to compute the triangulation matrix information necessary for estimating the 3D coordinates. These calibration steps were carried out using the open-source computer vision library (OpenCV) methods `OpenCV.findChessboardCorners`, `OpenCV.cornerSubPix`, `OpenCV.calibrateCamera`, and `OpenCV.stereoCalibrate` [32]. The videos were then trimmed by the user-specified trial onset and offset times to contain each utterance.

#### SPIGA

After initial data preprocessing, SPIGA placed markers on the lips, roughly around the vermillion border with a considerable amount of variability in its marker placement (see SPIGA in Fig 3). Thus, we averaged SPIGA’s marker coordinates across the first five frames of each video. In other words, we calculated the average lip coordinates across the first five frames. In addition, SPIGA’s upper lip coordinates were often slightly lower than the vermilion border. This inaccuracy diminished CoTracker’s performance, likely because the border provides a crucial cue for CoTracker. Therefore, as an additional correction step before CoTracker, an edge detection model using OpenCV (`OpenCV.Canny`) was ran to adjust upper lip coordinates to align with the vermilion border.

#### CoTracker

Due to the high computation power needed for CoTracker’s dense tracking capabilities, CoTracker was designed to process a smaller subsection of the original video that contained the subject’s face, where the crop is determined based on which segment of the video SPIGA identifies the face to be in on the first frame. The videos were cropped to a 704×512 resolution, which we found to be effective for both reducing computation power and allowing enough detail to show the entire face with extra padding in case of movement. Additional steps were taken at the end of the script to convert the tracking points from the dimension of the cropped video the tracking model ran on to the dimension of the original video, which was necessary for comparing the results with other tracking methods.

The notion of joint correlation also means that selecting supporting points, points designated to be tracked aside from the points of interest, has a large influence on the accuracy of the model’s output. We tested various types of supporting points, such as global dense grids, local dense grids, local points based on the lip contour, points from non-lip SPIGA landmarks, and combinations of these options. Consistent with the developers’ findings, our pilot data demonstrated that local supporting points (*i*.*e*., based on the contour/shape of the lips) combined with global supporting points yielded better performance than local or global points alone [30]. Hence, all data in the current study was generated by CoTracker configured with both local (*i*.*e*., points on the lips determined by SPIGA, see Fig 1) and global supporting points (*i*.*e*., a grid of points across the face and the background).

#### Postprocessing

After CoTracker identified pixel-wise coordinates from both the left and right videos, we triangulated points of interest in 3D space in a real-world unit (mm). During this process, the projection matrices were calculated using the camera calibration matrices that were computed in the preprocessing step. In addition, the OpenCV triangulation method, which relies on Direct Linear Transformation [45], was applied.

### Inpainting

We developed an inpainting pipeline built on a pretrained open-source Hugging Face stable diffusion inpainting model (stable-diffusion-v1-5 [39]). Because this model was trained using 512 *×* 512 pixel size images, we first identified the coordinate of a subsection of the video images (*i*.*e*., 512 *×* 512 pixel size) that contained the participant’s face and saved the cropped images from the videos using FFmpeg [33]. All of the images were in saved in the Portable Network Graphic (PNG) format to minimize the loss of image data.

An open software segmentation package from Meta (Segment Anything 2 [38]) was used to generate segmented binary masks. Following SAM2’s video predictor documentation, we calibrated their video segmentation model using the `sam2.1 heira large.pt` model checkpoint and the `sam2.1.heira l.yaml` model configuration setup. We modified the SAM2’s mask objection detection function to pinpoint and select the regions for segmentation. Using this approach, we targeted and segmented the AG501 sensors and wires. Binary masks were then created based on the segmentation and were used as inputs for inpainting.

Following segmentation, we preprocessed the masks with mask blurring and inflation guidelines documented by the developers [39]. We applied a mask blurring factor of 0.5 and a mask inflation factor of 10 for the `stable-diffusion-v1-5 model`. In addition, a scale factor of 5, a strength factor of 0.99, and the number of inference steps of 50 were applied for the model, which was identical to the “default” setup included in the code. The number of generations was set to 3, which means that the model generated 3 inpainted images and averaged them to create the final image. The inpainting model also incorporates a user-input prompt to direct the generative process, and the prompt “face of a young person, 4k, 5.3k” and a negative prompt of “blurry, unclear, deformed” were used. The resulting inpainted images were combined to generate a video for CoTracker.

## Data analysis

The two points of interest were defined as the mid-sagittal upper and lower vermilion borders. For all data analyses, we computed lip aperture (LA), the 3D Euclidean distance between these two points of interest:

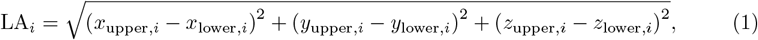

where *i* denotes each frame and *x, y, z* represents 3D axes defined by OpenCV’s 3D camera coordinate system. The subscripts, _upper_ and _lower_, represent coordinates at upper and lower lips respectively.

For precision tests, we simply computed the standard deviation for each video clip. All statistical tests were completed in R [46]. Using the lme4 package [47], two linear mixed-effects models were created, one for the minimal head movement dataset and another for the large head movement dataset. In each model, the tracking method (*i*.*e*., SPIGA vs. CoTracker) was included as a fixed effect, whereas participants was a random effect. We also included the video length (*i*.*e*. 1 vs. 0.5 s) as another fixed effect. The statistical significance (p-values) was determined using Type III analysis of variance via Satterthwaite’s method generated by the `lmerTest` package [48]. For post-hoc pairwise comparisons, we used the tukey method from the emmeans package [49].

For the accuracy test in which articulograph measurements were used as the “ground truth,” we first upsampled the CoTracker LA to 250 Hz to time-sync with the articulograph LA more precisely. We then used the maximum LA value in each LA data to coarsely align the two datasets temporally. After this initial alignment step, AG501 or CoTracker tracking was moved in an increment of a frame in both directions (*i*.*e*., moving the data to appear earlier in time or later in time), and the optimal time-synchronization alignment was defined as the alignment that yielded the lowest root mean square error (RMSE). In the current study, RMSE was defined as:

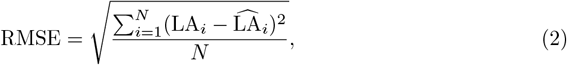

where LA_*i*_ and 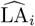 are lip apertures at a given frame *i*, measured by CoTracker and AG501 respectively. *N* represents the total sample size (*i*.*e*., the total number of frames).

After the temporal alignment step, CoTracker’s LA amplitude was normalized, because the locations where experimenters placed articulograph sensors were not identical to the locations where the points were placed by SPIGA (as part of the CoTracker pipeline). Hence, we first determined the local minimums in lip aperture (*i*.*e*., bilabial consonants) that correspond to the onset and offset of the target utterances (“Buy Bobby a puppy” or “puppy”, also see the shaded area in yellow in the right panels of Fig 5). By taking the averages of LA between the onset and offset (*i*.*e*., during the utterances), we normalized the CoTracker’s data to have the same average of the lip aperture of AG501 by adding or subtracting the discrepancy between the two averages. After these normalization steps, the final RMSE between CoTracker and AG501 was computed. All normalization steps and RMSE computation were completed with custom MATLAB scripts [50].

## Ethics Statement

Our study includes child participant. Our study procedures were approved by the Institutional Review Board at Purdue University (Protocol # IRB-2020-1025 and IRB-2023-744). All child participants provided verbal assent and their guardians or parents provided written informed consent prior to participating in our experiments.

## Supporting information

**Fig S1.**
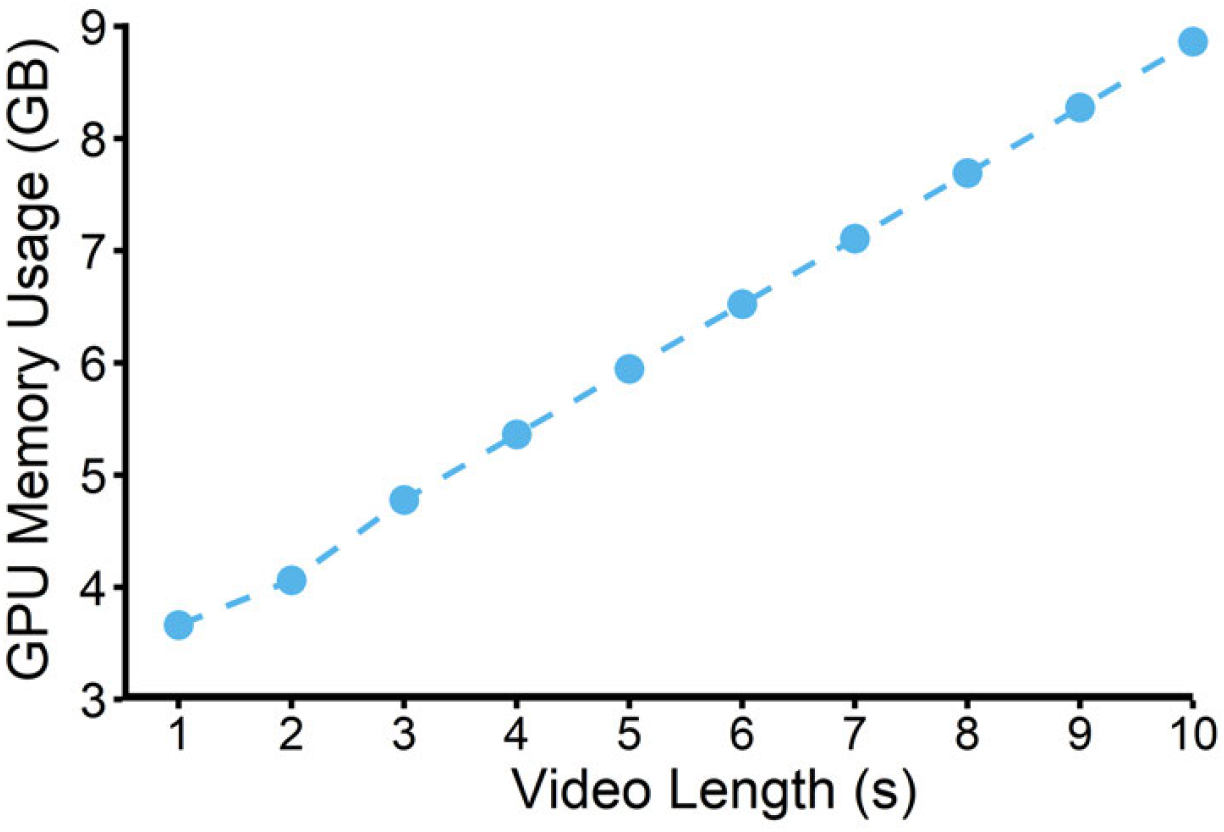

### GPU memory usage

CoTracker was used to extract lip landmarks from different time lengths of the same video (*i*.*e*., the first second, the first two seconds, and so on,) and the total amount of GPU memory usage on the computer (including the operation system processes) was recorded. The relationship between GPU memory and the video length was highly linear. Even for a 10 second-long video, the required GPU memory is around 10 GB.

## Acknowledgments

We thank Yuan Gao for conducting initial pilot tests and Kaelyn Loudermilk, Claney Outzen, and Phil Curtis for assisting with data collection.

## Funding

This research was funded by the National Institute of Health (R01DC018593 to A.B.).

## Notes

### Competing Interest Statement

The authors have declared no competing interest.

